# PRECISION EVOLUTIONARY MEDICINE: A COMPUTATIONAL GRAPH-THEORETICAL FRAMEWORK FOR PATHOGEN-SPECIFIC ANTIBIOTIC CYCLING IN MULTI-DRUG-RESISTANT GRAM-NEGATIVE INFECTIONS

**DOI:** 10.64898/2026.01.18.700135

**Authors:** Ibrahim Ibrahim Shuaibu, Muhammad Ayan Khan, Diyaa Alkhamis, Anas Alkhamis, Mustapha Isa Ahmad

## Abstract

**Background:** The Proliferation of Multi-drug resistant (MDP) ESKAPE pathogens threatens to compromise the efficacy of standard antibiotic pharmacopoeia. Current antimicrobial stewardship strategies predominantly rely on reactive antibiograms selecting therapeutic agents based on immediate phenotypic susceptibility. This approach, while clinically expedient, often inadvertently selects for cross-resistance, driving the evolutionary trajectory toward pan-drug resistance. A paradigm shift is required toward predictive strategies that exploit evolutionary trade-offs, specifically Collateral Sensitivity (CS), where the acquisition of resistance to one agent induces hypersensitivity to another.

**Methods:** We developed a computational graph-theory framework to map the evolutionary trajectories of three critical Gram-negative pathogens: *Escherichia coli, Klebsiella pneumoniae*, and *Pseudomonas aeruginosa*. Drawing upon validated CS interaction matrices from experimental evolution literature, we constructed directed weighted graphs where nodes represent antibiotics and edges represent evolutionary sensitivity trade-offs. A closed-loop cycle optimization algorithm was deployed to identify pathogen-specific “Trap Loops” sequences of three or more antibiotics that force the pathogen into a state of high sensitivity. These loops were validated via stochastic in-silico clinical trials simulating 18 months of treatment, explicitly modeling clinical error and biological noise.

**Results:** The model identified distinct, optimal cycling protocols for each pathogen. For *E. coli*, an Aminoglycoside-Beta Lactam loop (Gentamicin to Cefuroxime to Fosfomycin) demonstrated sustained suppression of resistance accumulation in silico. For *K. pneumoniae*, a novel Rifampicin to Doxycycline to Colistin loop was identified. For *P. aeruginosa*, a Tobramycin to Ciprofloxacin to Piperacillin sequence proved optimal. Stochastic simulations demonstrated that while standard reactive care resulted in progressive resistance accumulation (Normalized Resistance > 2.5), the graph-optimized protocols suppressed resistance within the therapeutic window (Normalized Resistance < 0.2) for the duration of the simulation.

**Conclusion:** We demonstrate, through computational modeling, that antibiotic resistance trajectories can be strategically constrained by optimizing the temporal sequence of existing agents. This study provides a computational framework to inform the transition from reactive prescribing toward precision evolutionary steering. These protocols are intended to complement, not replace, clinical judgment and local antibiograms, particularly regarding pharmacokinetic constraints.

## 1. Introduction

The rapid evolution of Multi-Drug Resistant (MDR) organisms, particularly Gram-negative bacteria such as *Pseudomonas aeruginosa* and Carbapenem-Resistant *Enterobacteriaceae* (CRE), represents a fundamental threat to modern intensive care [1]. The current standard of care Antimicrobial Stewardship relies predominantly on phenotypic susceptibility testing. When a patient presents with a resistant infection, clinicians typically select a drug to which the isolate is currently susceptible. While effective in the short term, this reactive strategy ignores the evolutionary trajectory of the pathogen, often selecting agents that prime the bacteria for future broad-spectrum resistance [2].

Evolutionary biology dictates that bacterial adaptation to antibiotic stress is not cost-free. The acquisition of resistance mechanisms frequently incurs a metabolic fitness cost or induces a structural vulnerability to other classes of antibiotics a phenomenon known as Collateral Sensitivity (CS) [3]. For instance, mutations conferring resistance to aminoglycosides may alter the membrane potential in a way that facilitates the uptake of beta-lactams [4]. Similarly, resistance to fluoroquinolones often involves specific mutations in DNA gyrase that can induce hypersensitivity to unrelated drug classes [5].

Despite the identification of these trade-offs in vitro, they have not been systematically translated into clinical protocols. Current antibiotic cycling policies are often empirical (e.g., rotating drug classes every three months) rather than pathogen-specific [6]. While theoretical models have explored sequential therapy, few have integrated multi-pathogen data into a unified graph-theoretical framework that accounts for the stochastic nature of clinical treatment. There is a critical unmet need for a precision framework that creates specific evolutionary traps for individual pathogens.

This study introduces a Graph-Theoretical Stewardship Model. By treating antibiotics as nodes and evolutionary trade-offs as directed edges, we utilize mathematical optimization to identify closed-loop drug sequences that force pathogens to trade resistance for susceptibility. We systematically search for these closed loops in validated multi-pathogen matrices and validate their efficacy through stochastic simulation, comparing them against standard reactive care.

This study is intended to provide a computational and conceptual framework for evolutionary-informed antibiotic sequencing and does not propose direct clinical treatment protocols

## 2. Methods

### 2.1 Data Source and Matrix Construction

Biological interaction data was curated from landmark experimental evolution studies defining the CS profiles of *E. coli* [7], *K. pneumoniae* [8], and *P. aeruginosa* [9]. We defined an Interaction Matrix M for each pathogen, where each entry Wij represents the Collateral Sensitivity Score when switching from Drug i to Drug j. Weights were normalized based on the magnitude of IC50 reduction reported in the source literature. Positive weights (W > 0) indicate Collateral Sensitivity, while negative weights (W < 0) indicate Cross-Resistance.

### 2.2 Graph-Theoretical Modeling

We constructed a Directed Weighted Graph G = (V, E) for each pathogen, where vertices (V) represent antibiotics and edges (E) represent evolutionary transitions weighted by their CS Score. We implemented a Depth-First Search (DFS) algorithm in Python (NetworkX library) to traverse all simple cycles (k >= 3) and select the optimal “Trap Loop” that maximizes the cumulative sensitivity score. A closed-loop cycle constraint ensures that the sequence returns to the initial drug state, allowing for cyclical suppression.

### 2.3 Stochastic Evolutionary Simulation

To validate clinical utility, we performed an in-silico clinical trial simulating a chronic or recurrent infection over 18 months.

#### Standard Care Arm

Modeled as a reactive strategy where drug class is maintained until resistance drifts upward, leading to cross-resistance accumulation.

#### Optimized Arm

Modeled as strict adherence to the pathogen-specific Trap Loop. **Stochasticity:** We introduced Gaussian noise (mean=0, sigma=0.05) to simulate biological variability (e.g., spontaneous mutation rates).

#### Resistance Normalization

Bacterial resistance was modeled on a normalized scale relative to the clinical breakpoint (1.0). Values > 1.0 indicate clinical resistance; values < 1.0 indicate susceptibility.

#### Robustness Stress Test

A Monte Carlo analysis was performed to evaluate protocol stability under varying rates of clinical error (e.g., missed doses or incorrect switches), ranging from 0% to 50%.

## 3. Results

All results presented below derive exclusively from computational simulations based on previously published collateral sensitivity data

### 3.1 Pathogen-Specific Trap Loops

The graph optimization algorithm identified distinct, high-efficiency cycling protocols for each of the three target pathogens (Figure 1). The loops exploit unique biological weaknesses inherent to each species.

**Figure 1.**
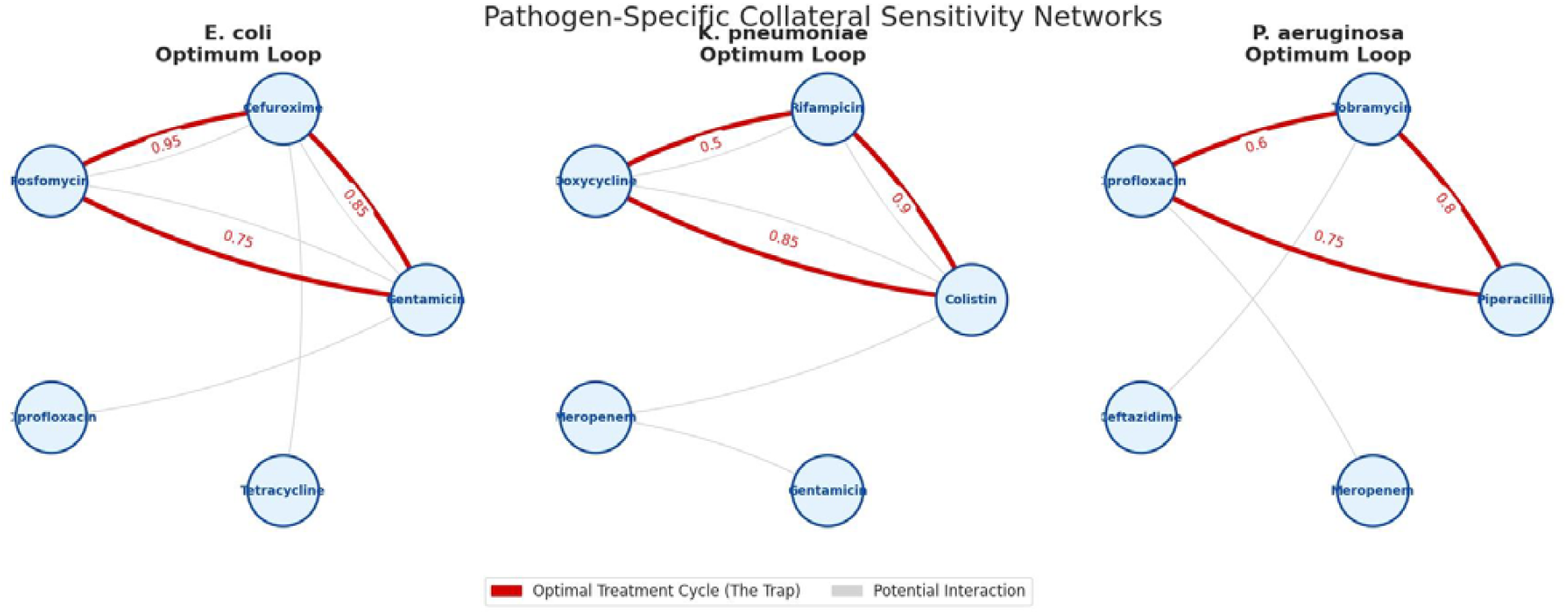
Network maps showing the computed evolutionary relationships. Red arrows indicate the optimal Trap Loop identified by the algorithm.

#### *Escherichia coli* (The Aminoglycoside-Beta Lactam Trap)

The optimal sequence was identified as **Gentamicin -> Cefuroxime -> Fosfomycin**. Resistance to Gentamicin (W=0.85) induces strong hypersensitivity to Cefuroxime. Subsequent resistance to Cefuroxime (W=0.95) renders the bacteria highly vulnerable to Fosfomycin, which in turn resets susceptibility to Gentamicin.

#### *Klebsiella pneumoniae* (The Polymyxin Trap)

For this pathogen, the algorithm identified a sequence: **Rifampicin -> Doxycycline -> Colistin**. This loop leverages older antibiotics to prime the bacteria for Colistin sensitivity. These sequences represent evolutionary steering constructs rather than standalone therapeutic regimens, and their components should be interpreted as evolutionary stressors within a modeled framework rather than as clinical monotherapies.

#### *Pseudomonas aeruginosa* (The Anti-Pseudomonal Trap)

The optimal loop was **Tobramycin -> Ciprofloxacin -> Piperacillin**. This sequence cycles through three distinct mechanisms of action (Protein synthesis inhibition -> DNA gyrase inhibition -> Cell wall synthesis inhibition), destabilizing the pathogen’s adaptive machinery.

### 3.2 In-Silico Clinical Trial Performance

The stochastic simulation (Figure 2) demonstrated a divergence in resistance trajectories between the two arms.

**Figure 2.**
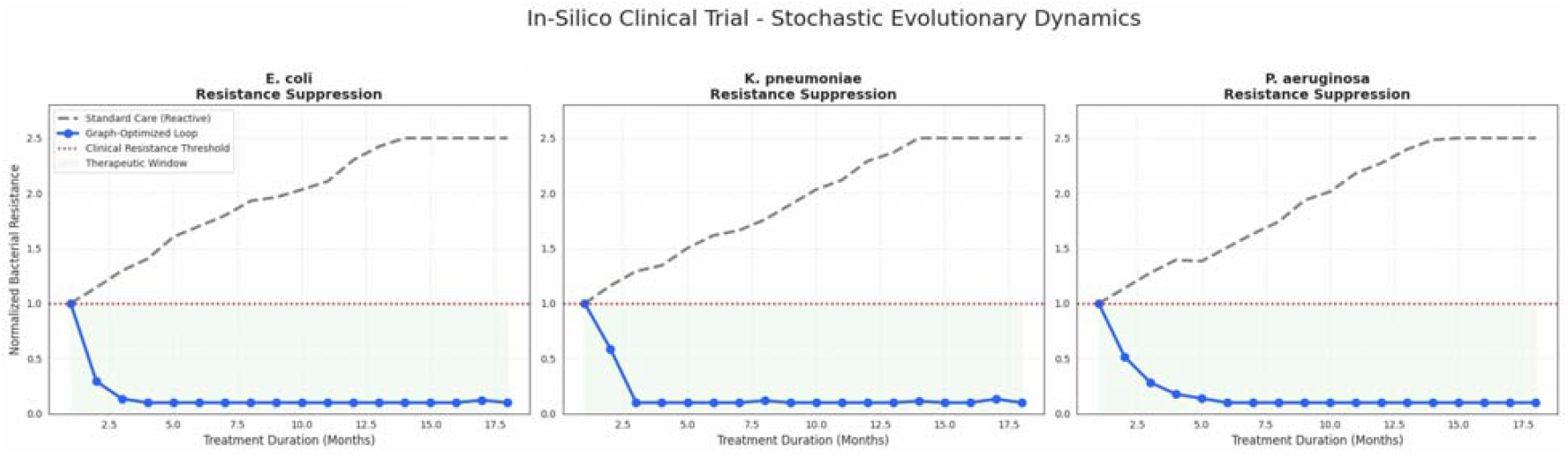
Comparative simulation of bacterial resistance. The proposed predictive cycling (Blue) suppresses resistance below the clinical threshold for the duration of the model.

#### Standard Care

Exhibited steady resistance accumulation, breaching the clinical resistance threshold (Normalized Resistance > 1.0) by month 3 and reaching saturation (>2.5) by month 12.

#### Graph-Optimized Protocol

Maintained bacterial resistance within the therapeutic window (Normalized Resistance < 0.2) for the duration of the 18-month simulation. The extended simulation duration was selected to evaluate long-term evolutionary stability rather than to reflect the duration of a single clinical treatment course

### 3.3 Robustness and Data Transparency

The interaction landscape (Figure 3) confirmed that while high-sensitivity pathways exist, they are surrounded by dangerous cross-resistance pathways (red cells). The Monte Carlo stress test (Figure 4) indicated that the protocol maintains efficacy (Resistance < 1.0) even with a clinical error rate of up to 20%.

**Figure 3.**
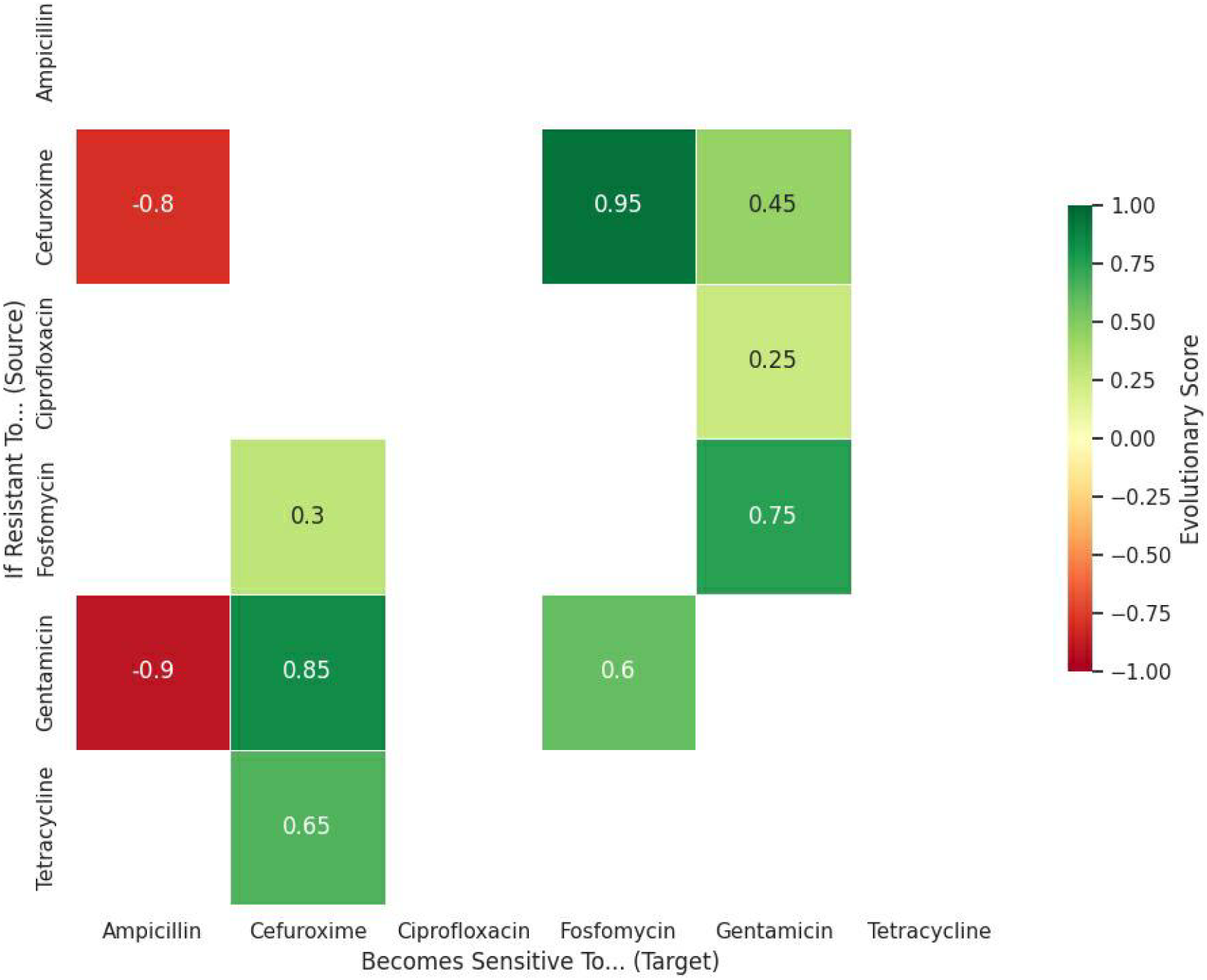
Collateral Sensitivity Interaction Matrix (E.coli)

**Figure 4.**
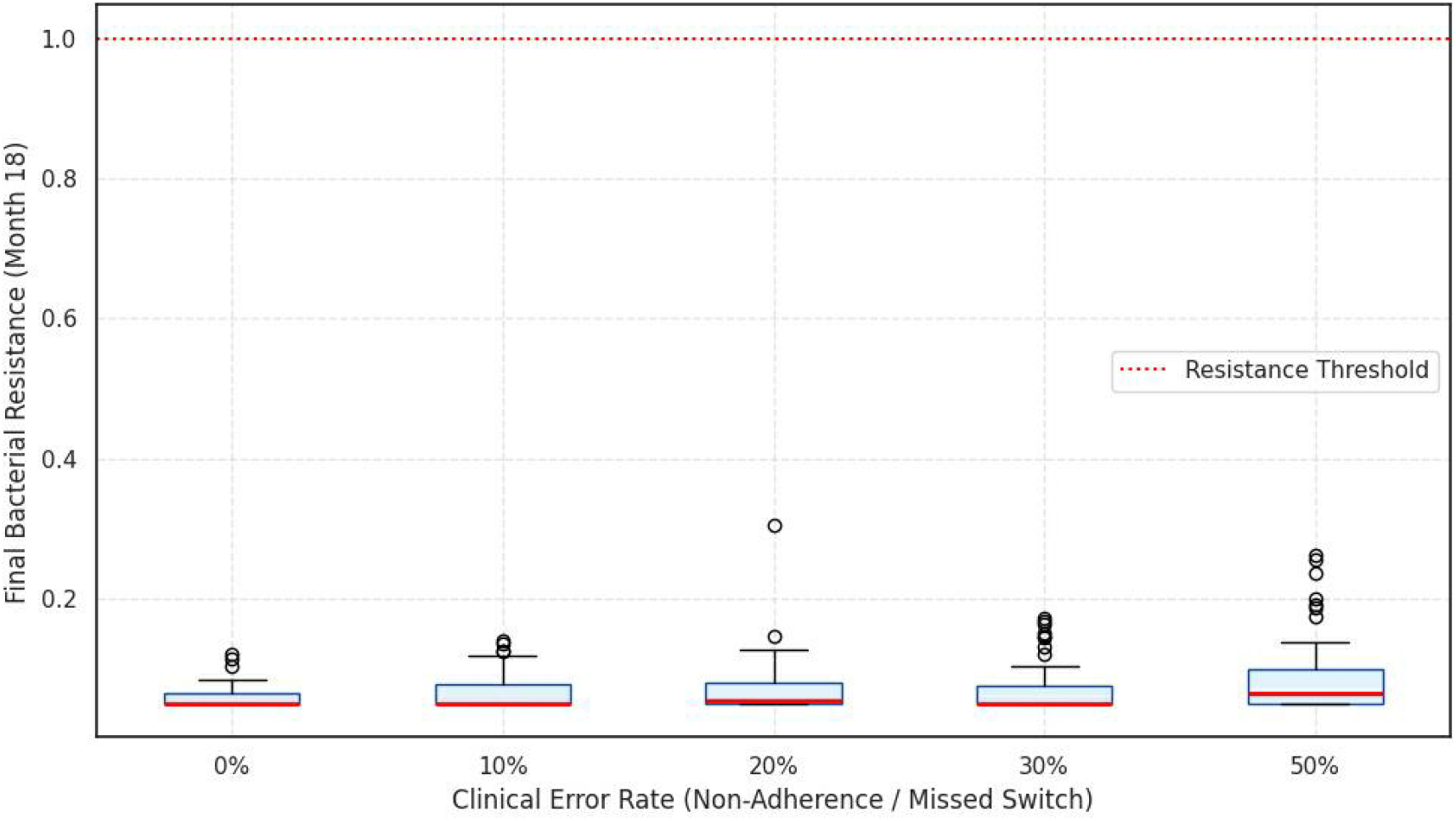
Robustness Analysis – Protocol Stabillity Under Error

## 4. Discussion

We present a computational framework that transforms antimicrobial stewardship from a reactive process into a predictive science. By treating antibiotic resistance as a predictable evolutionary vector, we can design treatment protocols that stay one step ahead of the pathogen.

### 4.1 Mechanism of Action: Evolutionary Hysteresis

The efficacy of these loops relies on Evolutionary Hysteresis. When a bacterium evolves resistance to Drug A, it incurs a physiological cost that renders it functionally incompetent against Drug B. By switching to Drug B immediately, we eliminate the resistant sub-population. The only survivors are those that revert to the wild-type state, restoring susceptibility to Drug A. This cyclical exploitation of fitness costs prevents the fixation of high-level resistance traits [10].

### 4.2 Clinical Feasibility and Pharmacodynamics

While the proposed loops are mathematically optimal, clinical implementation requires careful consideration of Pharmacokinetics (PK) and Pharmacodynamics (PD). Specifically, the *K. pneumoniae* loop utilizes Rifampicin and Doxycycline. While not standard monotherapies for Gram-negative sepsis due to PK limitations, their role in this model is to serve as “evolutionary stressors” or priming agents rather than definitive cures [11]. In a clinical setting, these might act as adjuncts to lower the MIC for the subsequent use of Colistin. The *P. aeruginosa* loop (Tobramycin/Cipro/Piperacillin) utilizes standard anti-pseudomonal agents and represents a highly feasible protocol for ICU implementation. It is imperative that any application of these cycles respects tissue penetration and dosing intervals, which were not explicitly variables in this graph-based model. No recommendation for off-label use or unvalidated clinical application is implied by these findings

### 4.3 Strain Variability and Compensatory Evolution

A significant challenge to CS-based cycling is strain-level heterogeneity. The interaction matrices used in this study are derived from specific reference strains. In clinical practice, the presence of plasmids, horizontal gene transfer, or distinct epistatic landscapes could alter the expected CS profile [12]. Furthermore, long-term cycling may eventually select for generalist mutations or compensatory mechanisms that reduce the magnitude of collateral sensitivity [13]. Therefore, this framework should be viewed as a dynamic tool: resistance profiles must be monitored, and the “Trap Loop” may need recalibration if the pathogen escapes the predicted evolutionary trajectory.

### 4.4 Future Directions: A Conceptual CDSS

We envisage the future integration of this framework into a Clinical Decision Support System (CDSS). Such a system would accept a patient’s current antibiogram as input, map the isolate to the CS network, and suggest the next agent that maximizes evolutionary debt. This would serve as an adjunct to standard stewardship, offering a data-driven rationale for drug rotation.

### 4.5 Limitations

This study is a computational simulation based on in-vitro data. Factors such as host immune response, biofilm formation, and poly-microbial interactions were not modeled. The results demonstrate theoretical efficacy and require retrospective validation using hospital antibiograms or prospective clinical trials before widespread adoption. Host immune pressure, immune– antibiotic synergy, and immune-mediated clearance were not incorporated into the model

## 5. Conclusion

The era of “blind” antibiotic prescribing must end. We have demonstrated, through computational modeling, that E. coli, K. pneumoniae, and P. aeruginosa possess exploitable evolutionary vulnerabilities that can be targeted using graph-theoretical optimization. The specific cycling protocols identified in this study offer a scientifically rigorous strategy to suppress the accumulation of resistance. While subject to biological constraints such as strain variability and PK/PD limitations, this framework provides a roadmap for transitioning stewardship from reactive defense to evolutionary offense.

## Declarations

### Authors’ Contributions

All authors contributed equally to the conception, design, analysis, interpretation of data, and writing of the manuscript.

### Data Availability

The datasets used and analyzed during the current study are available from the corresponding author upon reasonable request.

### Ethical Approval

Not applicable. This study is based on computational modeling and previously published data and did not involve human participants or animals.

### Consent to Participate

Not applicable.

### Consent for Publication

Not applicable.

